# Lipocalin 2 promotes inflammatory breast cancer tumorigenesis and skin invasion

**DOI:** 10.1101/2021.03.18.436052

**Authors:** Emilly S. Villodre, Xiaoding Hu, Richard Larson, Pascal Finetti, Kristen Gomez, Wintana Balema, Shane R. Stecklein, Ginette Santiago-Sanchez, Savitri Krishnamurthy, Juhee Song, Xiaoping Su, Naoto T. Ueno, Debu Tripathy, Steven Van Laere, Francois Bertucci, Pablo Vivas-Mejía, Wendy A. Woodward, Bisrat G. Debeb

**Affiliations:** Department of Breast Medical Oncology, The University of Texas MD Anderson Cancer Center, Houston, TX, USA; Department of Radiation Oncology, The University of Texas MD Anderson Cancer Center, Houston, TX, USA; Laboratory of Predictive Oncology, Aix-Marseille University, Inserm, CNRS, Institut Paoli-Calmettes, CRCM, Marseille, France; Department of Biological Sciences, The University of Texas at Brownsville, Brownsville, TX, USA; Department Biochemistry and Cancer Center, University of Puerto Rico Medical Sciences Campus San Juan, Puerto Rico; Department of Pathology, The University of Texas MD Anderson Cancer Center, Houston, TX, USA; Department of Biostatistics, The University of Texas MD Anderson Cancer Center, Houston, TX, USA; Department of Bioinformatics and Computational Biology, The University of Texas MD Anderson Cancer Center, Houston, TX, USA; Center for Oncological Research (CORE), Integrated Personalized and Precision Oncology Network (IPPON), University of Antwerp; MD Anderson Morgan Welch Inflammatory Breast Cancer Clinic and Research Program, The University of Texas MD Anderson Cancer Center, Houston, TX, USA

**Keywords:** lipocalin 2, LCN2, inflammatory breast cancer, skin invasion, brain metastasis

## Abstract

Inflammatory breast cancer (IBC) is an aggressive form of primary breast cancer characterized by rapid onset and high risk of metastasis and poor clinical outcomes. The biological basis for the aggressiveness of IBC is still not well understood and no IBC-specific targeted therapies exist. In this study we report that lipocalin 2 (LCN2), a small secreted glycoprotein belonging to the lipocalin superfamily, is expressed at significantly higher levels in IBC versus non-IBC tumors, independently of molecular subtype. LCN2 levels were also significantly higher in IBC cell lines and in their culture media than in non-IBC cell lines. High expression was associated with poor-prognosis features and shorter overall survival in IBC patients. Depletion of LCN2 in IBC cell lines reduced proliferation, colony formation, migration, and cancer stem cell populations in vitro, and inhibited tumor growth, skin invasion, and brain metastasis in mouse models of IBC. Analysis of our proteomics data showed reduced expression of proteins involved in cell cycle and DNA repair in LCN2-silenced IBC cells. Our findings support that LCN2 promotes IBC tumor aggressiveness and offer a new potential therapeutic target for IBC.

## 1. Introduction

Inflammatory breast cancer (IBC) is the most aggressive and deadly variant of primary breast cancer. Although IBC is considered rare in the United States (1%-4% of all breast cancer cases), it accounts for a disproportionate 10% of breast cancer-related deaths because of its aggressive proliferation and metastasis and limited therapeutic options [1-5]. IBC disproportionately affects young and African American women [1, 6]. IBC is associated with unique clinical and biological features and a distinctive pattern of recurrence with high incidence in central nervous system, lung, and liver as first site of relapse [4, 7, 8]. Even with multimodality treatment strategies, survival rates for women with IBC are far lower than for those with other types of breast carcinoma (non-IBC), with estimated 5-year overall survival rates limited to 40% versus 63% for non-IBC [4, 6-9]. These features underscore the critical need to better define the mechanisms that drive the aggressive behavior of IBC and to develop novel agents to improve the overall prognosis for women with IBC. Efforts have been undertaken to identify pathways and therapeutic targets distinct to IBC and to better elucidate the mechanisms of IBC aggressiveness [10-15]. However, the molecular and cellular basis for IBC aggressiveness remains unclear. Identification of specific targets and unraveling the mechanisms of growth and metastasis of this aggressive disease could lead to improvements in IBC patient survival.

Lipocalin 2 (LCN2, also known as neutrophil gelatinase-associated Lipocalin [NGAL], siderocalin, or 24p3) is a 25-kDa secreted glycoprotein that belongs to the lipocalin superfamily. LCN2 is known to sequester iron, as it binds siderophore-complexed ferric iron with high affinity, and has significant roles in immune and inflammatory responses, angiogenesis, cell proliferation, survival and resistance to anticancer therapies [16-21]. LCN2 has been implicated in the progression of several types of human tumors, including breast cancer, through several mechanisms, such as stabilization of MMP-9, sequestration of iron, induction of epithelial-mesenchymal transition, apoptosis resistance, lymphangiogenesis, and cell cycle arrest [16, 17, 19-26]. Moreover, high LCN2 expression levels have been linked with poorer survival in patients with breast cancer [17, 25, 27, 28]. Little is known regarding the oncogenic role of LCN2 in IBC tumors.

In the present study, we demonstrate that LCN2 was expressed at significantly higher levels in patients with IBC and that LCN2 promoted tumor growth, skin invasion, and metastasis in xenograft mouse models of IBC.

## 2. Materials and Methods

### 2.1. Cell lines

The SUM149 cell line was purchased from Asterand (Detroit, MI), and MDA-IBC3 cell line were generated in Dr. Woodward’s lab [29, 30], and cultured in Ham’s F-12 media supplemented with 10% fetal bovine serum (FBS) (GIBCO, Thermo Fisher, Carlsbad, CA), 1 µg/mL hydrocortisone (#H0888, Sigma-Aldrich, St. Louis, MO), 5 µg/mL insulin (#12585014, Thermo Fisher), and 1% antibiotic-antimycotic (#15240062, Thermo Fisher). HEK293T cells were purchased from the American Type Culture Collection (Manassas, VA, USA) and cultured in Dulbecco’s modified Eagle’s medium (DMEM) supplemented with 10% FBS and 1% penicillin and streptomycin (#15140122, Invitrogen, Carlsbad, CA, USA). All cell lines were kept at 37°C in a humidified incubator with 5% CO_2_ and were authenticated by short tandem repeat (STR) profiling at the Cytogenetics and Cell Authentication Core at UT MD Anderson Cancer Center.

### 2.2. Lentivirus-mediated knockdown

LCN2 stable knockdown clones were generated in SUM149 or MDA-IBC3 cells by using shRNA (shLCN2-1: TRCN0000060289 from Sigma-Aldrich; shLCN2-2: RHS4430-200252675 or shLCN2-3: RHS4430-200246537 from MD Anderson’s Functional Genomics Core Facility. The MISSION(R) pLKO.1-puro Empty Vector (SHC001, Sigma) was used as control (shCtl). HEK293T cells were transfected with 4.05 µg of target plasmid, pCMV-VSV-G (0.45 µg; #8584, Addgene;) and pCMV delta R8.2 (3.5 µg, #12263, Addgene) by using Lipofectamine 2000 (Life Technologies) for 24 h. SUM149 and MDA-IBC3 cells were incubated with the supernatant-containing virus plus 8 µg/mL of polybrene for 24 h. Stable cell lines were selected with 1 ug/mL of puromycin.

### 2.3. RNA isolation and real-time PCR

RNA was isolated by using TRIzol Reagent (Life Technologies) according to the manufacturer’s instructions. The cDNA was obtained with a High Capacity cDNA Reverse Transcription Kit with RNase Inhibitor (Thermo Fisher Scientific). Real-time PCR was done by using Power SYBR Green PCR Master Mix (Applied Biosystems) on a 7500 Real-Time PCR system (Applied Biosystems, Foster City, CA). LCN2 forward primer: 3’-CCACCTCAGACCTGATCCCA-5’, reverse primer: 3’-CCCCTGGAATTGGTTGTCCTG-5’; GAPDH forward primer: 3’-GAAGGTGAAGGTCGGAGT-5’, reverse primer: 3’-GAAGATGGTGATGGGATTTC-5’.

### 2.4. ELISA

Human Lipocalin-2/NGAL Quantikine ELISA Kits (#DLCN20, R&D Systems) were used to measure the levels of LCN2 in the cell lines according to the manufacturer’s instructions. Samples were assayed in duplicate.

### 2.5. Western blotting

Cells were lysed in RIPA buffer (Sigma) supplemented with 10 µL/mL phosphatase and 10 µL/mL protease inhibitor cocktail. SDS-PAGE and immunoblotting was carried out as described elsewhere [29]. The following primary antibodies were used: LCN2 antibody (1:1000, #MAB1757SP, R&D Systems, Minneapolis, MN, USA) or GAPDH (1:5000, #5174, Cell Signaling, Danvers, MA, USA) and samples were incubated overnight at 4°C. Secondary antibodies (1:5000), anti-rat IgG (#HAF005, R&D Systems) and anti-rabbit IgG (#7074, Cell Signaling), were incubated with the samples for 2 h at room temperature.

### 2.6. Proliferation

About 2,500 cells were seeded in triplicate in a 96-well plate. Cell proliferation was measured every day for up to 72 hours with the CellTiter-Blue assay (#G8080, Promega, Madison, WI) according to the manufacturer’s instructions. Absorbance was recorded at OD595 nm with a Multifunctional Reader VICTOR X 3 (PerkinElmer, Waltham, MA).

### 2.7. Colony-formation assay

About 100 SUM149 or 500 MDA-IBC3 shRNA Control or LCN2-silenced cells were plated in triplicate in 6-well plates. After 15 days, cells were fixed with methanol for 2 min, and stained with 0.2 % (w/v) crystal violet for 30 min. Colonies were counted by using GelCount (Oxford Optoronix, Abingdon, UK).

### 2.8. Migration and invasion assay

For the migration assay, 50,000 cells per well (triplicate) were seeded in medium without serum onto 8-μm polypropylene filter inserts in Boyden chambers (Fisher). Medium with 10% FBS was added onto the well. After 24 h, cells on the bottom of the filter were fixed and stained with Thermo Scientific Shandon Kwik Diff Stains (Fisher). The invasion assay was done as described above, except that the 8-μm polypropylene filter inserts were coated with Matrigel (#CB-40234, Corning, USA) and incubated for 24 h. Ten visual fields were randomly chosen under microscopy and cells were quantified by using ImageJ software (National Institutes of Health, Bethesda, MD, USA).

### 2.9. Mammosphere assay

For primary mammosphere formation, 30,000 SUM149 or MDA-IBC3 control or LCN2 knockdown cells were plated in ULTRALOW attachment 6-well plates (Corning, Inc.) in mammosphere medium (serum-free MEM supplemented with 20 ng/mL of bFGF [Gibco], 20 ng/mL epidermal growth factor [Gibco], B27 1x [Gibco], and gentamycin / penicillin / streptomycin [Thermo Fisher]). After 7 days, 5 ug/mL of MTT (Sigma-Aldrich) was added for 30 min and the mammospheres were counted by using GelCount (Oxford Optoronix). For secondary mammosphere formation, primary mammospheres were dissociated, counted, and 10,000 cells were plated in the ULTRALOW attachment 6-well plates in mammosphere media and analyzed after 7 days.

### 2.10. CD44/CD24 flow cytometry

About 2.5×10^5^ cells were suspended in CD24-PE mouse anti-human (#555428, BD Biosciences) or CD24-BV421 Mouse Anti-Human (#562789, BD Biosciences) and CD44-FITC mouse anti-human (#555478, BD Biosciences) or CD44-APC Mouse anti Human (#559942, BD Biosciences) solutions and incubated for 20 min on ice. Cells only, PE/BV421 only, and FITC/APC only were used as controls to set the gating. Fluorescence was detected by using a Gallios Flow Cytometer (Beckman Coulter, Brea, CA) at the Flow Cytometry and Cellular Imaging Core Facility (UT MD Anderson Cancer Center). FlowJo software (Treestar, Ashland, OR) was used to analyze the data.

### 2.11. Kinase Enrichment Analysis

The RPPA data was also used for the phosphoproteomic Analysis using kinase enrichment analysis (KEA - https://maayanlab.cloud/kea3/) [31]. Briefly, the 20 proteins that exhibit the highest phosphorylation fold change levels in control versus LCN2-silenced cells were analyzed. Two different analyses were performed using KEA: (1) the differentially phosphorylated proteins are queried for enrichment of kinase substrates; and (2) the differentially phosphorylated proteins are queried for enrichment of interacting proteins across 7 databases. The latter analysis is more general and is not limited to only kinase substrates. Both analyses result in the detection of kinases that are putatively responsible for the observed phosphorylation differences. Identified proteins by both analyses were mapped onto the STRING network (https://string-db.org) to investigate their mutual interactions.

### 2.12. In vivo experiments

Four- to six-week-old female athymic SCID/Beige mice were purchased from Harlan Laboratories (Indianapolis, IN). All animal experiments were done in accordance with protocols approved by the Institutional Animal Care and Use Committee of MD Anderson Cancer Center, and mice were euthanized when they met the institutional criteria for tumor size and overall health condition. For primary tumor growth, cells were injected into the orthotopic cleared mammary fat pad of mice as previously described [32]. Briefly, 5×10^5^ SUM149 shRNA Control / LCN2 knockdown cells were injected (9 mice / Control; 10 mice / LCN2 KD). Tumor volumes were assessed weekly by measuring palpable tumors with calipers. Volume (V) was determined as V = (L x W x W) x 0.5, with L being length and W width of the tumor. To determine latency, the first day when palpable tumors appeared was used to plot the graph. For brain metastatic colonization studies, we followed our lab protocol [33]. Briefly, 1×10^6^ MDA-IBC3 GFP-labeled shRNA Control / LCN2 knockdown cells (10 mice/group) were injected via the tail vein into SCID/Beige mice. At 12 weeks after tail-vein injection, mice were euthanized, and brain tissue collected and imaged with fluorescent stereomicroscopy (SMZ1500, Nikon Instruments, Melville, NY). ImageJ was used to measure GFP-positive areas to quantify the area of brain tumor burden. For mice with more than one brain metastasis, the area of each metastasis was considered and measured.

### 2.13. Statistical analysis

All in vitro experiments were repeated at least three times, and graphs depict mean ± SEM. Statistical significance was determined with Student’s *t* tests (unpaired, two-tailed) unless otherwise specified. One-way analysis of variance was used for multiple comparisons. Mann-Whitney test was used when normality was not met. *LCN2* expression in breast cancer samples was analyzed in the IBC Consortium dataset [34] for IBC and from a meta-dataset previously published [35]. Tumor samples were stratified as *LCN2*-high when expression in tumor was at least 2-fold the mean expression level measured in the normal breast samples; otherwise, the sample was classified as *LCN2*-low. Kaplan-Meier curves and log-rank tests were used to compare survival distributions. Univariate and multivariate Cox regression models were used to evaluate the significance of LCN2 expression on overall survival. A *p* value of <0.05 was considered significant. GraphPad software (GraphPad Prism 8, La Jolla, CA) was used.

## 3. Results

### 3.1. LCN2 mRNA is highly expressed in inflammatory breast cancer

Previous studies have shown that high LCN2 expression levels were correlated with poor prognosis in breast cancer patients [17, 25-27]. We further validated these findings by analyzing a meta-dataset of 8951 breast cancers, in which 87% of tumor samples were classified as *LCN2*-low and 13% as *LCN2*-high. Table 1 summarizes the clinico-pathological patient characteristics stratified by *LCN2* expression status. High expression of *LCN2* was associated with variables commonly associated with poor outcome: younger patients’ age, high grade, advanced stage tumors (pN-positive and pT3), ductal type, estrogen receptor (ER)-negative status, progesterone receptor (PR)-negative status, ERBB2-positive status, and aggressive molecular subtypes (ERBB2+ and triple-negative breast cancer [TNBC] subtypes). In this cohort, we also analyzed the association of *LCN2* expression and survival over time using the Kaplan–Meier method. We found that *LCN2*-high tumors had significantly shorter overall survival (*p*<0.0001) than *LCN2*-low tumors (Fig.1A).

**Table 1.** Clinico-pathological characteristics of tumor samples from patients with inflammatory breast cancer (IBC) or non-IBC according to *LCN2* expression.

| Covariate | Level | N | *LCN2*-low | *LCN2*-high | *p*-value |
| --- | --- | --- | --- | --- | --- |
| **Age (years)** | ≤50 | 2587 | 2218 (36%) | 369 (42%) | 1.10E-04 |
|  | >50 | 4520 | 4018 (64%) | 502 (58%) |  |
| **Pathological Grade** | 1 | 717 | 680 (13%) | 37 (4%) | <1.00E-06 |
|  | 2 | 2549 | 2359 (43%) | 190 (22%) |  |
|  | 3 | 3016 | 2389 (44%) | 627 (73%) |  |
| **Pathological Node (pN)** | Negative | 3666 | 3253 (57%) | 413 (53%) | 3.89E-02 |
|  | Positive | 2788 | 2426 (43%) | 362 (47%) |  |
| **Pathological Size (pT)** | pT1 | 2116 | 1912 (38%) | 204 (31%) | 2.00E-06 |
|  | pT2 | 2931 | 2588 (52%) | 343 (53%) |  |
|  | pT3 | 604 | 498 (10%) | 106 (16%) |  |
| **Pathological type** | Ductal | 4027 | 3492 (78%) | 535 (86%) | 3.00E-06 |
|  | Lobular | 500 | 471 (11%) | 29 (5%) |  |
|  | Other | 574 | 519 (12%) | 55 (9%) |  |
| **Estrogen Receptor status<sup>1</sup>** | Negative | 2753 | 1955 (25%) | 798 (71%) | 1.97E-215 |
|  | Positive | 6198 | 5875 (75%) | 323 (29%) |  |
| **Progesterone Receptor status<sup>1</sup>** | Negative | 4635 | 3746 (48%) | 889 (80%) | 3.06E-86 |
|  | Positive | 4284 | 4055 (52%) | 229 (20%) |  |
| **ERBB2 status<sup>1</sup>** | Negative | 7862 | 6975 (89%) | 887 (79%) | 2.37E-21 |
|  | Positive | 1089 | 855 (11%) | 234 (21%) |  |
| **Hormone Receptor subtype<sup>1</sup>** | HR+/ERBB2- | 5914 | 5598 (72%) | 316 (28%) | <1.00E-06 |
|  | ERBB2+ | 1089 | 855 (11%) | 234 (21%) |  |
|  | TNBC | 1938 | 1368 (17%) | 570 (51%) |  |
| **Overall Survival<sup>2</sup>** |  | 4984 | 1.00 | 1.58 [1.34 - 1.86]<sup>3</sup> | 3.31E-08 |

<sup>1</sup>mRNA status; <sup>2</sup>Univariate analysis; <sup>3</sup>hazard ratio (95% confidence interval)
Abbreviations: HR, hormone receptor; ERBB2, Erb-B2 Receptor Tyrosine Kinase 2 (HER2); TNBC, triple-negative breast cancer

Analysis of microarray data from the IBC World Consortium Dataset [34] consisting of IBC and non-IBC patient samples (n=389; IBC=137, non-IBC=252) showed that *LCN2* expression was significantly higher in tumors from IBC patients compared to non-IBC (*p*=0.0003; Fig 1B). We validated this finding in another independent data set [36] that compared mRNA expression of micro dissected IBC and non-IBC tumors (*p*=0.0379; Fig 1C). Here too, *LCN2* expression was higher in ER-negative IBC patients compared to ER-positive (*p*=0.0009; Fig. 1D) and in more aggressive subtypes, ERBB2-positive and TNBC, compared to hormone receptor (HR)-positive/ERBB2-negative subtype (Fig. 1E). Multivariate analysis showed that *LCN2* was expressed significantly higher in IBC tumors relative to non-IBC tumors, independently from the molecular subtype differences (Odds ratio, 1.71, *p*=0.034, Table 2). Here too, the survival analysis in IBC patients showed that *LCN2*-high tumors had significantly shorter overall survival (*p*=0.0317) than *LCN2*-low tumors (Fig.1F). Consistent with the patient data, the levels of *LCN2* were higher in IBC cell lines (Fig. 1G) and in the supernatants collected from IBC cell lines relative to non-IBC (Fig. 1H).

**Table 2.** Univariate and multivariate Cox regression analysis of inflammatory breast cancer (IBC) patient samples versus non-IBC (n=389).

| IBC vs. non-IBC | Univariate |  |  | Multivariate |  |  |
| --- | --- | --- | --- | --- | --- | --- |
|  | Odds-ratio | 95% CI | p-value | Odds-ratio | 95% CI | p-value |
| LCN2, high vs low | 2.09 | 1.43 - 3.06 | 1.43E-03 | 1.71 | 1.13 - 2.6 | 3.42E-02 |
| Molecular subtype |  |  |  |  |  |  |
| ERBB2+ vs HR+/ERBB2- | 2.82 | 1.82 - 4.38 | 1.02E-04 | 2.5 | 1.59 - 3.93 | 8.16E-04 |
| TNBC vs HR+/ERBB2- | 1.9 | 1.22 - 2.97 | 1.69E-02 | 1.51 | 0.93 - 2.44 | 0.162 |

Abbreviations: HR, hormone receptor; ERBB2, Erb-B2 Receptor Tyrosine Kinase 2 (HER2); TNBC, triple-negative breast cancer

**Fig 1.**
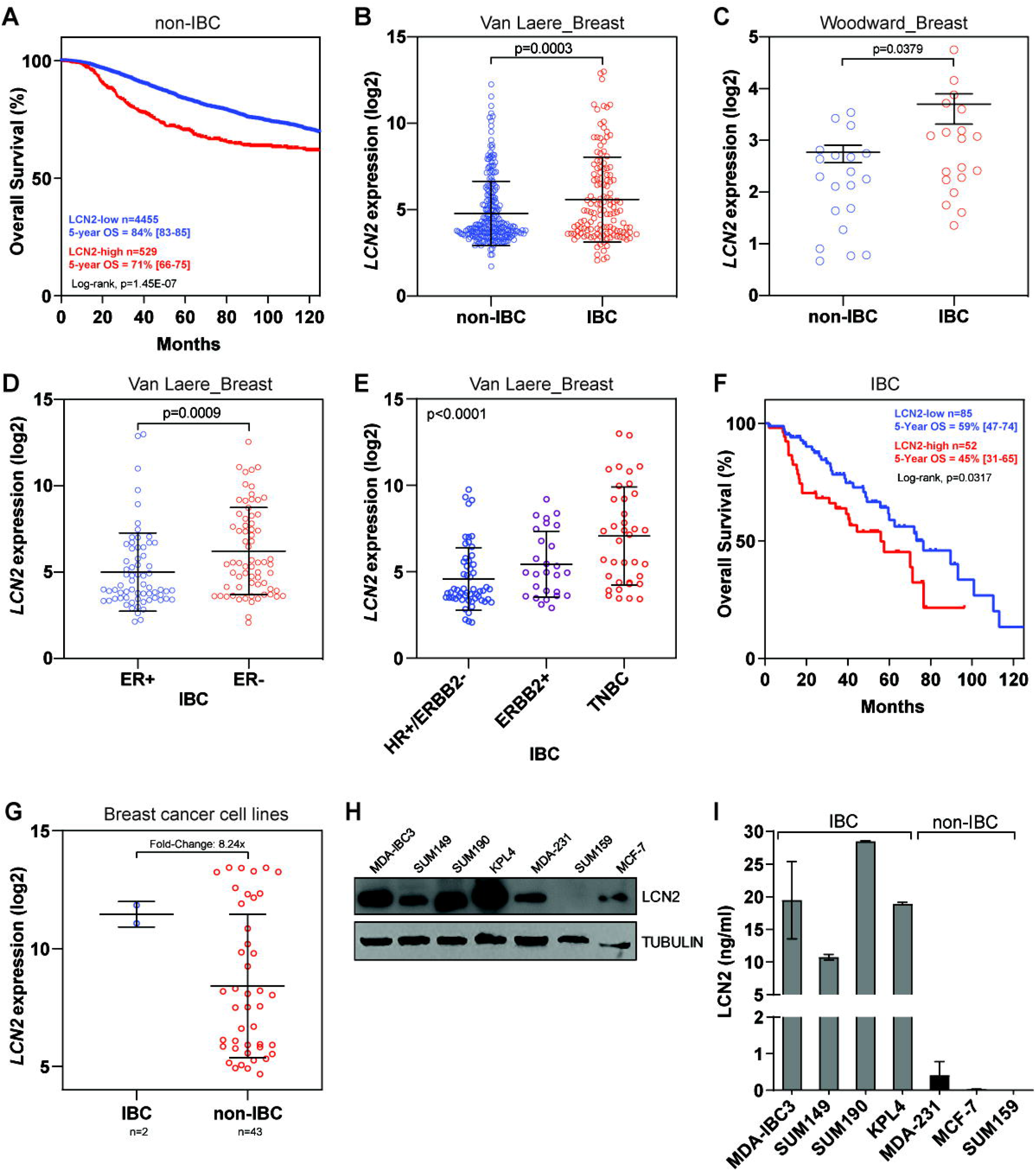
LCN2 was highly expressed in tumors from patients with IBC. **(A)** High *LCN2* expression was associated with shorter overall survival in a meta-dataset of patients with non-IBC. **(B-C)** *LCN2* mRNA expression was higher in tumors from IBC patients versus non-IBC patients in two independent breast cancer datasets [34, 36]. **(D)** *LCN2* mRNA expression was higher in estrogen receptor (ER)-negative compared to ER+ samples IBC samples. **(E)** *LCN2* mRNA expression was higher in more aggressive molecular subtypes, ERBB2+ and triple-negative breast cancer (TNBC), compared to hormone receptor (HR)-positive/HERBB2-negative subtype. **(F)** *LCN2*-high expression correlates with shorter overall survival in patients with IBC. **(G)** *LCN2* mRNA expression was higher in IBC cell lines compared to non-IBC cell lines. **(H-I)** LCN2 protein expression was higher in IBC cell lines compared to non-IBC cell lines shown by **(H)** immunoblotting or **(I)** ELISA for secreted LCN2 in supernatants. Graphpad Prism software was used to obtain the *p* values, with Mann-Whitney tests used to compare two categories or one-way analysis of variance to compare three or more categories.

Taken together, our findings show that LCN2 is highly expressed in IBC tumors and is correlated with aggressive features and poor outcome suggesting it may contribute to the aggressive pathobiology of IBC tumors.

### 3.2. LCN2 knockdown reduced aggressiveness features in vitro

We generated stable LCN2 knockdown cell lines [SUM149 (triple-negative IBC); MDA-IBC3 (HER2+ IBC)] to investigate the role of LCN2 in IBC aggressiveness in vitro and in vivo. LCN2 knockdown was confirmed by qRT-PCR and immunoblotting (Fig. 2A,B). Because LCN2 is a secreted protein, we evaluated levels of LCN2 protein in the supernatants from control and LCN2-silenced IBC cell lines by using ELISA. We observed significant reduction of secreted LCN2 in the LCN2-silenced IBC cells (Fig. 2C). Silencing LCN2 slightly reduced proliferation of SUM149 cells but did not affect MDA-IBC3 cells (Fig 2D). Depletion of LCN2 reduced the capacity of the cells to form colonies (Fig. 2E) and to migrate and invade (Fig. 3A,B). LCN2 silencing also significantly reduced the percentage of cancer stem cell populations in LCN2-silenced IBC cells relative to control, as shown by reductions in primary and secondary mammosphere formation efficiency (Fig. 3C,D) and CD44^+^CD24^-^ cell subpopulations (Fig. 3E). These findings indicate that suppression of LCN2 in IBC cells reduced in vitro aggressiveness features.

**Fig 2.**
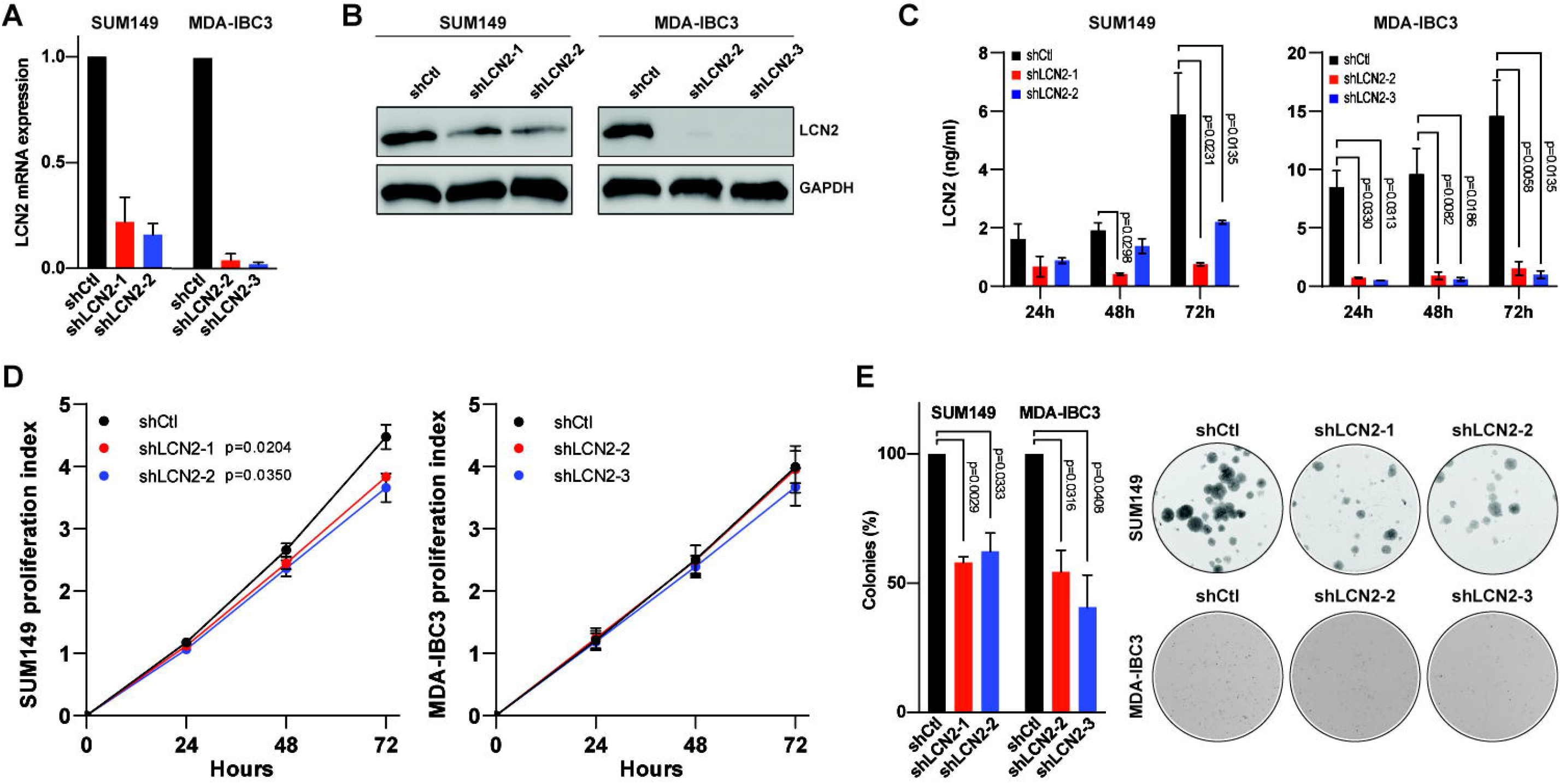
Silencing LCN2 decreased colony formation efficiency. LCN2 was knocked down (shLCN2) in two IBC cell lines (SUM149 and MDA-IBC3) and confirmed by **(A)** qRT-PCR and **(B)** immunoblotting. **(C)** Secreted LCN2 measured in control and silenced cells by ELISA at the indicated times. **(D)** Proliferation was evaluated in control and LCN2-silenced SUM149 and MDA-IBC3 cells with CellTiterBlue assay on the indicated days. **(E)** Cells were seeded in low numbers to measure the capacity to form colonies in LCN2 knockdown and control.

**Fig 3.**
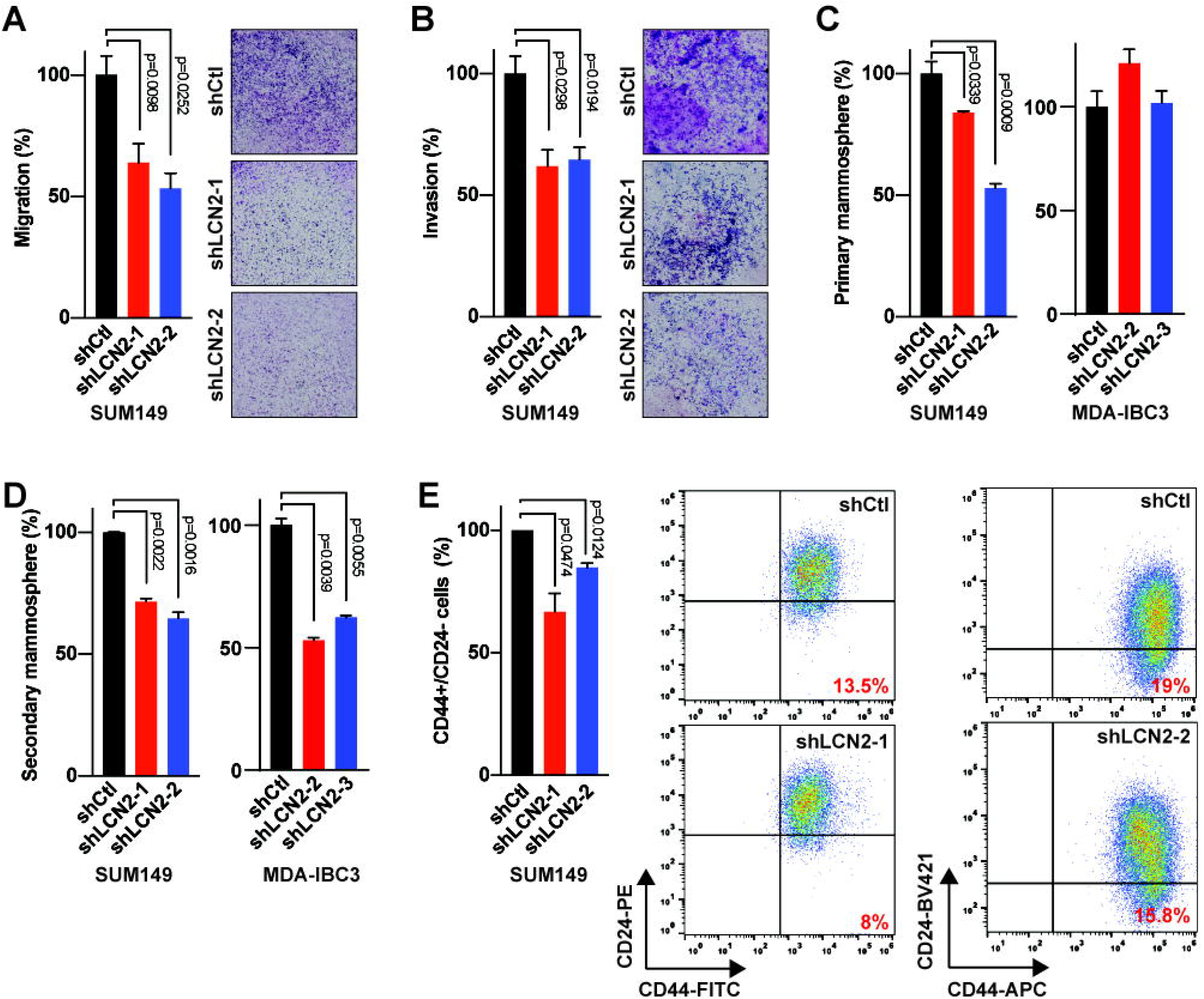
LCN2 knockdown reduced aggressiveness features in vitro. **(A)** Migration and **(B)** invasion by control cells (shCtl) and LCN2-knockdown (shLCN2) SUM149 cells. **(C)** Primary mammosphere formation efficiency and **(D)** secondary mammosphere formation efficiency. **(E)** CD44^+^CD24^-^ cells (marker of cancer stem cells) were measured by flow cytometry.

### 3.3. Silencing of LCN2 inhibited tumor growth and skin inavsion

To investigate the effects of LCN2 on tumor growth and skin invasion, key characteristics of IBC tumors [4], we injected SUM149 control or LCN2-silenced cells into the cleared mammary fat pad of SCID/Beige mice. Silencing of LCN2 reduced tumor volumes (*p*=0.0037; Fig. 4A) and tumor latency, *i*.*e*. the ability to initiate tumor growth: mice transplanted with SUM149 LCN2-silenced cells took longer to initiate tumors than did those transplanted with SUM149 control cells (*p*=0.0145, Fig. 4B). Because IBC typically manifests with skin invasion and formation of tumor emboli [4], we assessed skin invasion visually during primary tumor growth, as evidenced by loss of fur at the tumor site and skin redness and thickness, and during tumor excision when tumors were firmly connected with the skin. Analysis of resected tumors showed that significantly fewer mice with SUM149 LCN2-silenced cells had skin invasion/recurrence compared with mice implanted with control cells (shLCN2: 2 of 8 mice [25%] vs. shControl: 7 of 8 mice [87.5%], *p*=0.01; Fig. 4C;4D). On histologic examination, tumors generated from LCN2-silenced cells were more differentiated than those generated from control SUM149 cells (Fig. 4E); we further observed tumor emboli, another hallmark of IBC tumors, in SUM149 control-transplanted tumors but not in tumors generated from LCN2-silenced SUM149 cells (Fig. 4E).

**Fig 4.**
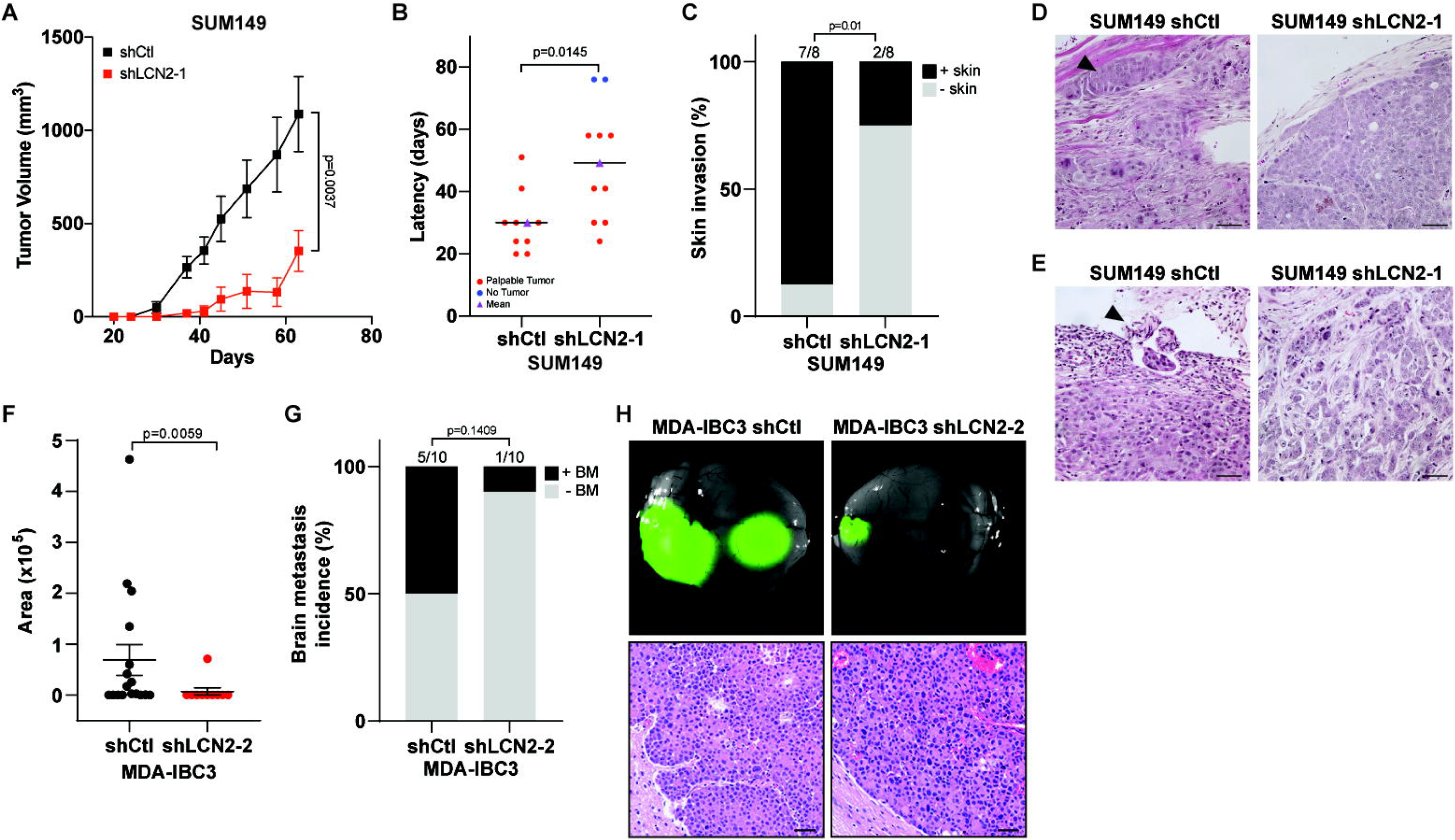
Silencing LCN2 inhibited tumor growth and skin inavsion. **(A-C)** SUM149 shRNA Ctl or LCN2-knockdown (shLCN2) cells were transplanted orthotopically into the cleared mammary fat pad of SCID/Beige mice (n= 9/Ctl; 10/shLCN2) and tumor volume measured weekly; **(A)** tumor volume, **(B)** tumor latency, and **(C)** incidence of skin invasion/recurrence after resection of primary tumors. **(D-E)** Hematoxylin and eosin staining of primary tumors generated from LCN2 control and knockdown SUM149 cells. Both **(D)** skin invasion and **(E)** tumor emboli, two hallmarks of IBC, appeared only in the control-derived tumors (arrow head). Scale bar, 100 µm. **(F)** Metastatic burden (area) of each brain metastasis formed was quantified by using ImageJ software. BM, brain metastasis. **(G)** Incidence of brain metastasis. N=10 mice per group. Fisher’s exact test was used to obtain *p* values. **(H)** Top, green fluorescent protein (GFP) imaging of brain metastasis lesions generated from tail-vein injection of GFP-labeled MDA-IBC3 shRNA Ctl or LCN2 knockdown cells, and bottom, hematoxylin and eosin stains of brain metastasis lesions. Scale bar, 50 µm.

We recently generated xenograft mouse models of brain and lung metastasis via tail-vein injection of IBC cell lines [29, 33]. We also showed that sublines of SUM149 generated from brain metastases (BrMS) and lung metastases (LuMS) have distinct morphologic and molecular features [29]. Microarray profiling of these sublines showed upregulation of *LCN2* in the brain metastatic sublines (Supplementary Fig. S1A), and we confirmed higher levels of secreted LCN2 in the BrMS sublines versus LuMS by ELISA (Supplementary Fig. S1B). Most recently, Chi et al elegantly demonstrated that LCN2 promotes brain metastatic growth in mouse models of leptomeningeal metastasis, highlighting a potential brain metastasis-promoting role for LCN2 [37]. We investigated the functional role of LCN2 in IBC brain metastasis by using our HER2+ MDA-IBC3 mouse model, which has a high propensity to metastasize to the brain and has been used to identify targets and develop therapeutics against brain metastasis [29, 38-40]. We found that the brain metastatic burden was significantly lower in mice that had received tail-vein injection of LCN2-silenced MDA-IBC3 cells than in mice injected with control cells (Fig. 4F, *p*=0.0059). Also, fewer mice injected with LCN2-silenced cells developed brain metastasis (1 of 10 [10%]) than did mice injected with control cells (5 of 10 mice [50%]), although this trend was not statistically significant (*p*=0.1409; Fig. 4G). Representative stereofluorescence and hematoxylin and eosin images of brain metastases are shown in Fig. 4H. Overall, our findings suggest that LCN2 may drive IBC tumor progression, skin invasion/recurrence, and brain metastasis.

### 3.4. LCN2 silencing impairs cell cycle-associated proteins

To identify potential mechanisms and pathways involved in suppression of tumor growth and skin invasion in LCN2-silenced cells, we used reverse phase proteomics assay (RPPA) profiling to compare control and LCN2-silenced SUM149 cells. Our analysis showed reduced expression of cell cycle-associated proteins (such as AXL, FOXM1, Chk1, CDK1, Wee1, Aurora-B, and cyclin-B1 and the mTOR/AKT pathway) in LCN2-silenced IBC cells (Fig. 5A). Gene set enrichment analysis revealed several key signaling pathways that were enriched in the control cells, including those associated with cell cycling, DNA repair and mTOR signaling (Fig. 5B). Furthermore, we performed kinase enrichment analysis (KEA) [31] on the 20 proteins that exhibited the highest phosphorylation fold changes in LCN2-control vs. LCN2-silenced SUM149 cells (Supplementary Table 1). Based on the set of predicted activated kinases (Supplementary Table 1 and Supplementary Table 2), an interaction network was generated (Fig. 5C). Based on the node degree distribution (i.e. the distribution of the number of interactions per gene in the network), MAPK1 (N=10), MAPK8 (N=7), RPS6KB1 (N=7) and MTOR (N=11) appear to be central to LCN2 action in SUM149 cells. Thus, LCN2 may regulate different pathways, including cell cycle and mTOR proteins to promote tumor growth in IBC.

**Fig 5.**
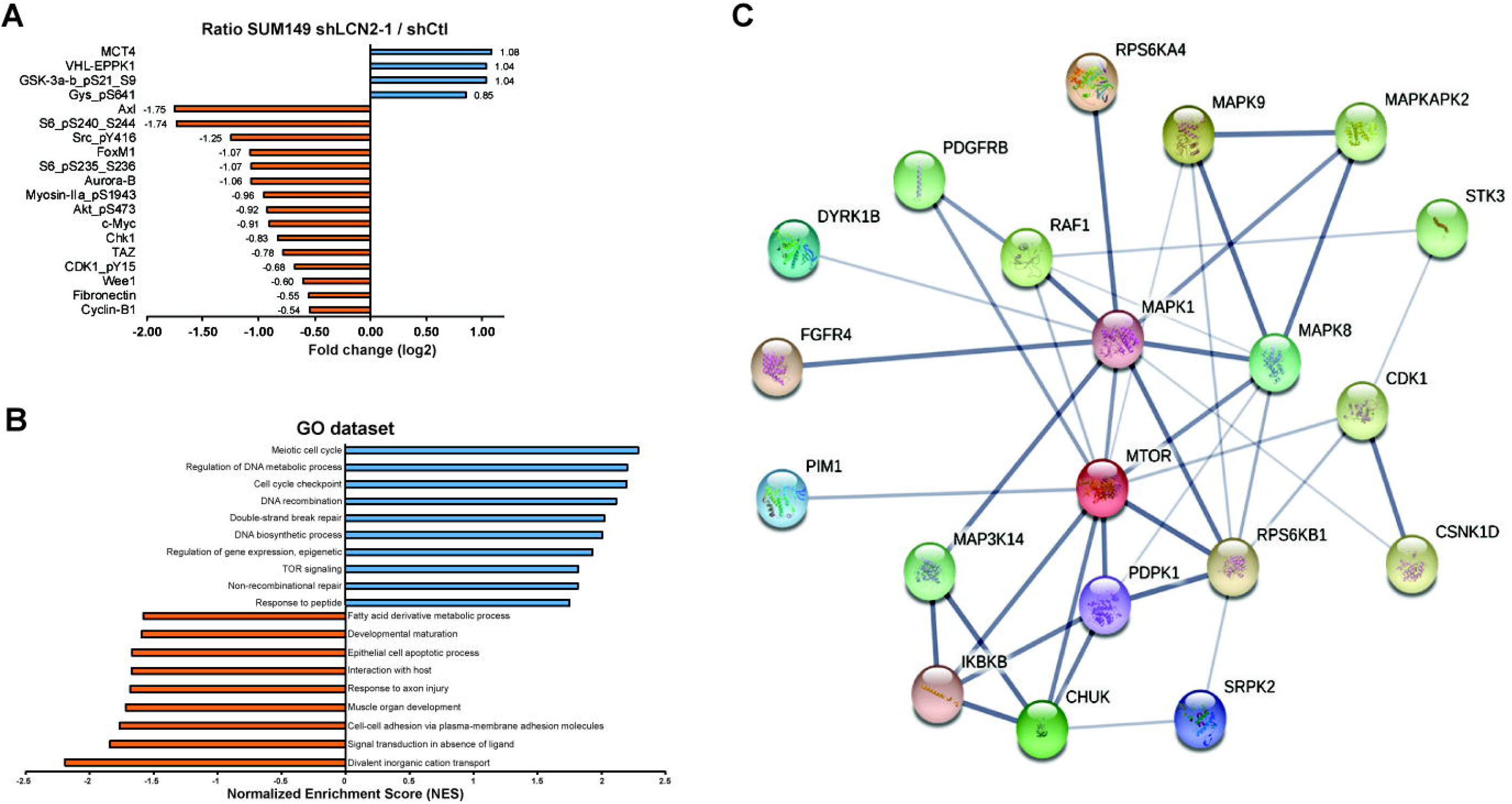
Silencing of LCN2 impairs cell cycle-associated proteins. **(A)** The top proteins downregulated in LCN2-silenced cells compared with control cells after reverse phase protein array (RPPA) proteomic analysis. **(B)** Gene set enrichment analysis of RPPA data identified pathways that are enriched or downregulated in control vs. LCN2-silenced SUM149 cells. **(C)** STRING interaction network of predicted active kinases based on enrichment of kinase substrates and protein interactions identified using kinase enrichment analysis. The confidence of the interaction is reflected by the edge thickness. Based on node distribution analysis, four central proteins were identified (MAPK1, MAPK8, RPS6KB1 and MTOR).

## 4. Discussion

Inflammatory breast cancer is an aggressive form of breast cancer with poor survival outcomes. Although considerable effort has been undertaken to understand the unique biology of IBC, insights are still limited as to the molecular properties that mediate the development and aggressiveness of IBC. Herein, we report that the secreted glycoprotein LCN2 was highly expressed in tumors from IBC patients and in IBC cell lines. We further demonstrate, with in vitro and in vivo studies, that LCN2 has a tumor promoter function in IBC.

LCN2 has been implicated in the progression of several types of human tumors. LCN2 expression is higher in solid tumors than in corresponding normal tissues [24, 41], and it is mainly described as tumor promoter in many cancers, including pancreas, glioblastoma, thyroid, kidney, esophagus, and breast cancer [20, 28, 42-48].

In breast cancer, increased LCN2 expression was associated with poor outcomes and shown to be an independent prognostic marker of disease-specific-free survival [27, 48, 49]. LCN2 also correlates with several important unfavorable prognostic factors in breast cancer, such as hormone-negative status, high proliferation levels, high histologic grade, and the presence of lymph node metastases [27, 48, 49]. Further, serum levels of LCN2 have been shown to correlate with cancer progression and higher likelihood of metastasis in breast cancer [25, 50]. The oncogenic role of LCN2 has been reported in xenograft and LCN2-knockout mouse models. Disruption of the *LCN2* gene in MMTV-PyMT mice was found to suppress primary tumor formation without affecting lung metastasis [51]. Using the spontaneous MMTV-*ErbB2*(V664E) LCN2^-/-^ mouse model, Leng et al reported delayed tumor growth and reduced lung metastasis burden in these LCN2^-/-^ mice [16]. Another group showed that injection of wild-type PyMT tumor cells into LCN2-deficient mice did not alter primary tumor formation but did significantly reduce lung metastasis [52]. LCN2 has also been shown to promote tumor progression in xenograft mouse models [17, 25]. Consistent with these studies, our current work with xenograft mouse models of IBC supports that LCN2 has a tumor promoter function in IBC tumors. We demonstrated that silencing of LCN2 reduced tumor initiation and growth, skin invasion/recurrence, and brain metastasis burden in preclinical mouse models of IBC.

We further reported that depletion of LCN2 in IBC cell cultures reduced features associated with aggressiveness in vitro, including migration, invasion, and cancer stem cell populations. Others have also found that reduction of LCN2 levels affected the same features in MDA-MB-231 cells (triple-negative breast cancer cell line) and in SK-BR-3 (HER2+ breast cancer cell line) [16, 25]. However, our data demonstrating higher levels of secreted LCN2 in IBC versus non-IBC cell lines and showing significant inhibition of key IBC tumor features such as tumor emboli/skin invasion in LCN2-silenced tumors suggest that LCN2 may exert its influence via an IBC-specific mechanism. The LCN2 protein has many functions, including transport of fatty acids and iron, induction of apoptosis, suppression of bacterial growth, and modulation of inflammatory responses [16, 17, 19-21, 25, 53]. In malignant cells, LCN2 promotes oncogenesis through several mechanisms, including stabilization of MMP-9, sequestration of iron, induction of EMT, apoptosis resistance, and regulation of cell cycling [16, 17, 19-21, 25, 53]. Here we report that LCN2 could regulate cell cycle-associated proteins such as FOXM1, Chk1, CDK1, Aurora-B, Wee1, and cyclin-B1 to promote its oncogenic role in IBC tumors. Others have also found that silencing of LCN2 affected the expression of cell cycle proteins by reducing cyclin-D1 and inducing p21, resulting in G0-G1 cell cycle arrest [22-24].

LCN2 is also a potential therapeutic target in cancer and other diseases. An antibody against LCN2 was found to decreased lung metastasis in a 4T1-induced aggressive mammary tumor model [16]. In cervical cancer cells, treatment with LCN2-neutralizing antibody reduced the migration and invasion of cells that overexpressed LCN2 [54]. In other diseases, use of an anti-LCN2 neutralizing antibody showed reductions in reperfusion injury after stroke and attenuated skin lesions in a psoriasis mouse model [55, 56]. These findings suggest that LCN2 could be an exploitable therapeutic target in IBC and other aggressive tumors. Further studies are needed to explore therapeutic strategies in IBC models by using antibodies against LCN2 or targeting LCN2-associated molecular pathways, including those involved in cell cycling.

In summary our studies provide evidence, for the first time, that LCN2 is highly upregulated in IBC tumors and that it is required for tumor growth and skin invasion in mouse models of IBC; our findings further suggest that LCN2 could be a therapeutic target for IBC and other aggressive cancers.

## Declaration of competing interest

We have no conflicts of interest to declare.

## Acknowledgements

We thank Christine F. Wogan, MS, ELS, of MD Anderson’s Division of Radiation Oncology for scientific editing and review of the manuscript. The Functional Genomics Core Facility at UT MD Anderson Cancer Center and the Flow Cytometry and Cellular Imaging Core Facility are both funded through NCI grant P30 CA016672 to the University of Texas MD Anderson Cancer Center.

## Funding

This study was supported in part by the following grants: UPR/MDACC Partnership for Excellence in Cancer Research (U54CA096297-CA096300 to BGD), Susan G. Komen Career Catalyst Research Grant (CCR16377813 to BGD), and State of Texas Morgan Welch Welch Inflammatory Breast Cancer Program.

## Contributions

ESV and BGD conceived and designed the project, performed most of the experiments, analyzed the data, and interpreted the results. XH, RL, WB, SRS, and KG performed some experiments. PF, FB, CI, and XS helped with data analysis. JS provided statistical analysis support. SK provided pathological expertise and analysis of xenograft tumors. SVL, FB, GS, PV-M, NTU, WAW, and DT provided resources and contributed to revision of the manuscript. ESV and BGD wrote and edited the manuscript with input from all other authors.

## Figure Legends

**Additional file 1. Figure S1**. LCN2 expression is higher in sublines generated from brain metastasis (BrMS) than those generated from lung metastasis (LuMS). **(A)** Microarray analysis of sublines generated from BrMS or LuMS of SUM149 cells showed *LCN2* to be one of the top upregulated genes in BrMS (red arrow). Samples are described in Debeb 2016 [29]. **(B)** LCN2 is secreted in higher levels in BrMS versus LuMS.

**Additional file 2. Supplementary Table 1**. Top kinases predicted to be activated based on kinase-substrate interactions of differentially phosphorylated proteins.

**Additional file 3. Supplementary Table 2**. Top kinases predicted to be activated based on kinase-substrate and protein-protein interaction analysis of differentially phosphorylated proteins across 10 different knowledge bases.

## Notes

### Competing Interest Statement

The authors have declared no competing interest.

