## Supplementary figures and images for "Lipocalin 2 promotes inflammatory breast cancer tumorigenesis and skin invasion"

### Supplementary Figure 1

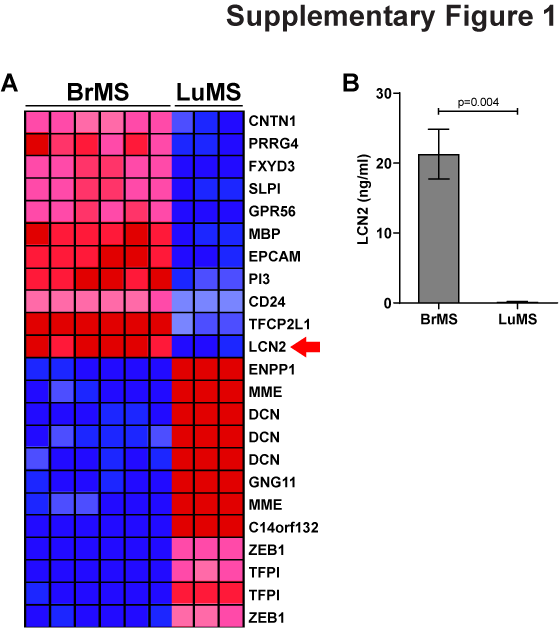
