## Supplementary Table 1 for "Lipocalin 2 promotes inflammatory breast cancer tumorigenesis and skin invasion"

**Supplementary Table 1**
Top kinases predicted to be activated based on kinase-substrate interactions of differentially phosphorylated proteins.

| Rank | Protein | Overlapping Protein | FET *p*-value | *FDR* | Odd Ratio |
|---|---|---|---|---|---|
| **1** | CSNK1D | 11 / 359 | 2.72E-12 | 9.21E-10 | 31.58 |
| **2** | MAP3K14 | 8 / 136 | 9.75E-12 | 1.65E-9 | 62.44 |
| **3** | CHUK | 13 / 773 | 6.83E-11 | 7.72E-9 | 17.09 |
| **4** | PDPK1 | 13 / 819 | 1.36E-10 | 1.15E-8 | 16.11 |
| **5** | IKBKB | 9 / 343 | 7.38E-10 | 5.00E-8 | 26.92 |
| **6** | RPS6KA4 | 8 / 256 | 1.37E-9 | 7.75E-8 | 32.49 |
| **7** | FGFR4 | 10 / 511 | 1.62E-9 | 7.85E-8 | 19.94 |
| **8** | STK3 | 7 / 180 | 3.05E-9 | 1.29E-7 | 40.42 |
| **9** | SRPK2 | 9 / 414 | 3.72E-9 | 1.40E-7 | 22.2 |
| **10** | DYRK1B | 9 / 437 | 5.9E-9 | 2.00E-7 | 21.01 |
