## Supplementary Table 2 for "Lipocalin 2 promotes inflammatory breast cancer tumorigenesis and skin invasion"

Top kinases predicted to be activated based on kinase-substrate and protein-protein interaction analysis of differentially phosphorylated proteins across 10 different knowledge bases.

| Rank | Protein | Mean rank | Overlapping Proteins |
|---|---|---|---|
| **1** | RPS6KB1 | 34.55 | 18 |
| **2** | MAPK8 | 37.91 | 19 |
| **3** | PDGFRB | 39.27 | 16 |
| **4** | MAPK9 | 42.1 | 17 |
| **5** | MAPKAPK2 | 46.78 | 15 |
| **6** | MAPK1 | 53.18 | 20 |
| **7** | RAF1 | 56.64 | 17 |
| **8** | CDK1 | 59.1 | 19 |
| **9** | MTOR | 59.45 | 18 |
| **10** | PIM1 | 60.5 | 15 |
